# Neuroprotective effects of exogenous erythropoietin in Wistar rats by downregulating apoptotic factors to attenuate N-methyl-D-aspartate-mediated retinal ganglion cells death

**DOI:** 10.1101/773754

**Authors:** Wen-Sheng Cheng, I-Hung Lin, Kathy Ming Feng, Zhi-Yang Chang, Yu Chuan Huang, Da-Wen Lu

## Abstract

The aim of this study was to investigate whether exogenous erythropoietin (EPO) administration attenuates N-methyl-D-aspartate (NMDA)-mediated excitotoxic retinal damage in Wistar rats. The survival rate of retinal ganglion cells (RGCs) were investigated by flat mount analysis and flow cytometry. A group of male Wistar rats were randomly assigned to five groups: negative control, NMDA80 (i.e., 80 nmoles NDMA intravitreally injected), NMDA80 + 10ng EPO, NMDA80 + 50ng EPO, and NMDA80 + 250ng EPO. The NMDA80 + 50ng EPO treatment group was used to evaluate various administrated points (pre-/co-/post-administration of NMDA80). Meanwhile, the transferase dUTP Nick-End Labeling (TUNEL) assay of RGCs, the inner plexiform layer (IPL) thickness and the apoptotic signal transduction pathways of μ-calpain, Bax, and caspase 9 were assessed simultaneously using an immunohistochemical method (IHC). When EPO was co-administered with NMDA, attenuated cell death occurred through the downregulation of the apoptotic indicators: μ-calpain was activated first (peak at ∼18hrs), followed by Bax and caspase 9 (peak at ∼40hrs). Furthermore, the morphology of RGCs has clearly demonstrated the visual recovery of IPL thickness at 40 hours after injection. Exogenous EPO successfully protected RGCs by downregulating apoptotic factors to attenuate NMDA-mediated excitotoxic retinal damage.

## Introduction

Glaucoma is one of the major causes of irreversible blindness worldwide [1, 2]. It is a group of optic neuropathies characterized by the loss of retinal ganglion cells (RGCs) [3]. Even though elevated intraocular pressure (IOP) is often a main indicator of glaucoma, it can also occur with normal IOP levels [4]. Several mechanisms may be responsible for RGC death, including apoptosis [5, 6], trophic factor withdrawal (TFW) [7, 8], inflammation [9, 10], and excitotoxicity [11]. Loss of the ganglion cell inner plexiform layer (IPL) is highly correlated with overall loss of visual field and is therefore a potential biomarker to evaluate glaucoma progression in patients [12]. Given the variety of conditions that could lead to RGC death, neuroprotection may be used to prevent the loss of IOP-independent RGC.

Glutamate, one of the common excitatory neurotransmitters in the retina, has long been known to exert excitotoxic actions on neurons of the inner retina [13]. The effects of glutamate on cells are mediated by ionotropic receptors that are classified into α-amino-3-hydroxy-5-methyl-4-isoxazolepropionic acid (AMPA), N-methyl-D-aspartate (NMDA) subtypes, or kainate receptors according to their preferred agonist [14]. NMDA receptors are activated by the co-agonists NMDA (or glutamate) and glycine, which are known to be predominantly involved in neuronal cell death in the retina and brain [15, 16]. In several studies, glutamate was shown to be involved in several retinal diseases including glaucoma [17], retinal ischemia [18, 19], and optic neuropathy [20].

Erythropoietin (EPO), a hematopoietic factor, has been confirmed to stimulate the differentiation and proliferation of erythroid progenitor cells. A few studies found that EPO and its receptors (EPOR) were expressed in retinal and brain tissues [21–23] and inhibits apoptotic activity in erythrocyte progenitors [24]. EPO was shown to have neuroprotective functions against light-induced retinal degeneration [25] and retinal ischemia in various studies on neuronal cell death [26]. Furthermore, there is some evidence that it may also be neuroprotective against human stroke [27–29]. In our previous study, EPO was protective of RGC against NMDA-, TNF-α-, and TFW-induced damage [30]. After EPO binds on EPOR, homodimerization begins and subsequently activates various signal transduction cascades, involving kinases such as RAS/RAF/ERK or the PI-3K/Akt kinase pathways [31]. The net effect of erythroblasts is enhanced differentiation and proliferation, inhibiting apoptosis [32–35].

Recently, new studies have demonstrated that EPO exerts neuroprotective effects against excitotoxin- and NO-induced apoptosis, which was shown in the crosstalk between the Jak2 and NF-κB signaling pathways [36]. Results from *in vitro* studies showed that a significant EPO signaling overlap appeared between erythroid progenitors and neuronal cells. EPO-mediated neuroprotection was observed against the excitotoxic and hypoxic deaths of cultured motoneurons, C19 teratoma cells, and hippocampal neurons. These findings further demonstrated that EPO display neuroprotective effects after mechanical trauma (e.g., retinal, cerebral, or spinal cord ischemia) or neuroinflammation [25,28,35,36]. Thus, the neuroprotective function of EPO has mostly been investigated under ischemic, hypoxic, and excitotoxic paradigms that involve cell death as a result of apoptosis and necrosis. Apoptosis activates several caspases in extrinsic and intrinsic pathways. The death-inducing signaling complex is activated in the extrinsic pathway leading to the recruitment of caspases whereas mitochondrial membrane depolarization was observed in the intrinsic pathway. Bax and Bak are pro-apoptotic proteins that control the outer mitochondrial membrane permeability [37]. When the membrane is depolarized, cytochrome c is released into the cytoplasm and forms the apoptosome complex with Apaf-1 (apoptotic protease activating factor-1) in the presence of ATP. Procaspases 9 is recruited and activated by the apoptosome leading to caspase activation downstream. These activated caspases subsequently cause DNA fragmentation and apoptosis. The calcium-activated neural protease (µ-calpain) is thought to be activated by the influx of intracellular calcium via the endoplasmic reticulum stress response pathway. Once µ-calpain is activated, downstream pro-apoptotic proteins may be activated and there may be crosstalk between caspases [38]. Recently, EPO was demonstrated to suppress the activation of caspases or the appearance of transferase dUTP Nick-End Labeling (TUNEL)-positive cells. It can even prevent mitochondrial depolarization. However, there were very few reports on the preventative effects of EPO on RGC death caused by glaucoma or chronic neurodegenerative diseases [39]. A previous study evaluated the effects of EPO on RGCs in episcleral vessel cautery-induced rats with glaucoma using 200ng EPO, which found no significant decrease in the RGC of the EPO-treated group [40–42]. However, little is known about how EPO signal transduction occurs *in vivo* and the practical usefulness of EPO in the prevention of chronic purely apoptotic neuronal cell death, which contributes to vision loss in glaucoma and the progression of neurodegenerative diseases.

The objective of this study was to establish NMDA-mediated excitotoxic retinal damage in Wistar rats, and then to determine the time course of neuroprotection of RGC by EPO and to determine the thickness of the segmented inner plexiform layer. In addition, the downregulation of the apoptotic signal transduction pathways of μ-calpain, Bax, and caspase 9 may involve the crosstalk in NMDA-mediated neurotoxic rats with EPO treatment.

## Materials and Methods

### Animal model

Male Wistar rats (Taiwan National Laboratory Animal Center, Taipei, Taiwan) weighing between 225 and 250 g, were kept in a temperature-controlled environment (21–22℃) under a 12-h light-dark cycle. All studies were conducted in accordance with the guidelines of the Association for Research in Vision and Ophthalmology Statement on the Use of Animals in Ophthalmic and Vision Research. The study protocol was approved by the Institutional Animal Care and Use Committee of National Defense Medical Center (Permit number: IACUC-07-175 and IACUC-08-209).

### Drug treatment

NMDA (Sigma, St. Louis, MO, USA)-mediated retinal neurotoxicity was established as reported previously by Nivison-Smith *et al* [19]. The rats were briefly anesthetized with a mixture of ketamine hydrochloride (50 mg/kg, Nang Kuang Pharmaceutical, Tainan, Taiwan, Republic of China) and xylazine hydrochloride (13.3 mg/kg, Sigma, St. Louis, MO, USA). After the topical administration of 0.5% proparacaine hydrochloride (Alcon Lab, Fort Worth, TX, USA), an intravitreal injection of 2 μL 40 mM NMDA in BSS PLUS solution (Alcon Lab, Fort Worth, TX, USA) was placed into the right eye of each rat using a 30-gage needle (Hamilton, Reno, NV, USA). The solution was administered into the sclera approximately 1 mm behind the limbus. The BSS PLUS solution was used as a vehicle control and the NMDA and different doses of EPO were used in the therapeutic treatment groups. Two of them were injected into the left eye.

### Retrograde labeling of retinal ganglion cells

RGCs were labeled with FG (Fluorogold) (Sigma, St. Louis, MO, USA) by injecting the FG solution into the superior colliculi using a stereotaxic device (Stoelting, Wood Dale, IL, USA) as described previously [43]. Seven days after NMDA induction, the rats were quickly anesthetized with the ketamine/xylazine mixture, and the skin over the cranium was then incised to expose the scalp. Two vertical holes 1 mm in diameter were drilled on both sides of the skull with a dentist drill 6 mm posterior to the bregma and 1.5 mm lateral to the midline. Two microliters of 3% FG solution was delivered using a micropipette 3.8, 4.0, and 4.2 mm beneath the bone surface.

### Retinal flat mount imaging and FG-labeled RGC counting

Seven days after the NMDA induction, the rats were euthanized with carbon dioxide (CO2) and the eyes were immediately enucleated. The retinas were dissected and fixed in 4% paraformaldehyde (Sigma, St. Louis, MO, USA) for 1 hour. After washing with phosphate buffered saline (PBS), the retinal flat mount samples were placed as four-radial incisions and were prepared on slides in PBS with 10% glycerol (Sigma, St. Louis, MO, USA). The retinal slides were examined under a fluorescence microscope (Olympus BX-50, Olympus Optical, Tokyo, Japan) using UV excitation (330–385 nm) and a barrier filter (420 nm) to determine the RGC cell number. Digital images were taken using a CCD camera (SPOT, Diagnostic Instruments, Sterling Heights, MI, USA). Each retina was visually divided into four quadrants (i.e., superior, inferior, nasal and temporal). The quadrants were further divided into central (0.8–1.2 mm from the optic disk), middle (1.8–2.2 mm from the optic disk), and peripheral regions (0.8–1.2 mm from the retinal border). Within each region, the cells falling in two fields (200 × 200 μm^2^ in size) were counted. Both manual and automatic counts were used to count RGCs. For automatic counts, the digital images were processed using Image J (http://rsbweb.nih.gov/nih-image/, U.S. National Institutes of Health, Bethesda, MD, USA) and the RGB (red-green-blue) images were converted to 8-bit grayscale for binary counting [22]. The RGCs were classified into three groups based on soma sizes described previously (small 9.4 μm, medium 9.4–12.6 μm, and large 12.6 μm) [30]. FG-labeled cells were counted and the mean number of RGCs per square millimeter was calculated.

### Flow cytometry

Retinal cells were evaluated using a flow cytometer (Calibur; Becton Dickinson, San Jose, CA, USA). The parameters of the cytometer were optimized with the following settings, which were based on preliminary studies: the FSC (forward scatter) value used for reflecting cell volume/size and the SSC (side scatter) value used for reflecting fluorescence intensity and its internal complexity. FL1, FL2, and FL3 reflect the green (FITC, fluorescein), orange (PE, phycoerythrin-R), and red (peridinin-chlorophyl protein (PerCP), peridinin-chlorophyl) labeled fluorescence intensities, respectively. These) were applied with a high flow rate (60 µl/min). The compensation values were determined from CaliBRITE three-color kit (BD Biosciences, Franklin Lakes, NJ, USA). The cell sizes were estimated from the FSC using a calibration curve, which was established by 6, 10, 15, and 20 μm diameters of polystyrene microspheres (Polysciences, Warrington, PA, USA). Ten-thousand cells per sample were counted automatically; and incubated cells without the fluorescence-conjugated antibody were used as an unstained blank. The expression of FG intensity and Thy1.1 were evaluated as geometric means of the FL2 and FL3 fluorescence intensities, respectively. The fluorescence intensity data was analyzed using CellQuest Pro (Becton Dickinson, San Jose, CA, USA) and FCS Express V3 software (De Novo Software, Los Angeles, CA, USA). The cell viability can be used to evaluate the health of cells due to the dye reacting with cellular amines. In necrotic cells, the reactive dye can enter the cell via the compromised membranes and react with the free amines in the interior and on the surface of the cell, resulting in an intense fluorescent staining in unhealthy or dead cells. Cell-surface staining of amines in the viable cells will result in relatively dim staining that can easily be distinguished from the bright staining of dead cells.

### Histochemical technique: TUNEL assay

TUNEL is a method used for detecting DNA fragmentation generated during apoptosis. It works by labeling the 3′ - hydroxyl termini in the double-strand DNA breaks. The dissected retinas were incubated in papain solution (composed of 20 U/mL papain, 1 mM L-cysteine, 0.5 mM EDTA, and 200 U/mL DNase I [Sigma, St. Louis, MO, USA] in Earle’s balanced salt solution [EBSS, Invitrogen, Carlsbad, CA, USA]) at 37℃ for 40 minutes. The retinas were transferred to ovomucoid-BSA buffer (1 mg/mL ovomucoid, 1 mg/mL BSA, and 100 U/mL DNase I in EBSS) for 5 minutes at 37℃. The tissues were then gently triturated through a plastic pipette until dispersion and the retinal cells were fixed with 2% paraformaldehyde in PBS for 20 minutes at room temperature. The cells were centrifuged and re-suspended in PBS containing 0.4% Triton X-100 (Sigma, St. Louis, MO, USA). 1% BSA was used as a nonspecific blocker. Each sample was subsequently centrifuged and re-suspended in DNA-labeling solution (APO-BrdU TUNEL Assay Kit; Invitrogen, Carlsbad, CA, USA) for 1 hour at 37℃ to label DNA strand breaks in the apoptotic cells. The cells were then incubated with a rabbit anti-Fluoro-Gold antibody (1:100, Millipore, Bedford, MA, USA) mouse PerCP-labeled anti-Thy1.1 antibody (1:100, BD Biosciences, Franklin Lakes, NJ, USA), and FITC(fluorescein)-labeled anti-BrdU antibody (1:200, eBioscience, San Diego, CA, USA) for 30 minutes. After washing with PBS, the cells were incubated with goat anti-rabbit IgG phycoerythrin-R (PE) conjugated antibody (1:200, Santa Cruz Biotechnology, Santa Cruz, CA, USA) for 30 minutes. The cells were washed with PBS and re-suspended in 1 mL PBS containing 0.1% Triton X-100. The total number of retinal cells was counted using a hemocytometer (Bright-Line, Reichert, Buffalo, NY, USA) and the size distributions of the cells were evaluated via flow cytometry.

### Histological analysis

After enucleation, the eyes were immersed overnight at 4°C in a fixative solution consisting of 4% paraformaldehyde in PBS (pH 7.4). This was followed by dehydration with gradient-increased ethanol (75- to 100%) and embedded in paraffin. The retina cross-section had a thickness of 4 μm was prepared and stained with hematoxylin and eosin. The morphology analysis was conducted using an optical microscope. The cell numbers of the ganglion cell layer (GCL) were counted 1.0 to 1.5 mm to the margin of the optic nerve head. Five samples were taken with an interval of 100 μm, and the mean thickness of the inner plexiform layer was analyzed. The thickness of the outer nuclear layer was used as the internal standard.

### Immunohistochemistry, IHC

The histosection of the retina was attached on gelatin-coated glass slides. The slides were dewaxed with xylene twice, rehydrated with a decreasing-gradient ethanol solution (100 to 80%), then rinsed three times with PBS. Citrate buffer (0.01M pH 6) was used to maintain boiling for 20 minutes and rinsed three times with PBS. BSA 0.1% was used as a blocking agent to prevent nonspecific binding of antigens and antibodies for 10 minutes. The primary antibodies of Bax (1:100), caspase-9 (1:100), and μ-calpain (1:100) were reacted for 1 hour individually. After washing with PBS, animal serum was used for a post-primary blocking agent for 15 mins. The Novolink polymer (anti-mouse/rabbit IgG-Poly-HRP) was reacted with the primary antibody for 15 minutes, 3, 3’-diaminobenzidine (DAB chromogen), and 0.02% hematoxylin to stain the nucleus. The coverslips were washed three times with PBS, the samples were embedded with Fluoromount-G and then examined under an optical microscope. GCL and positive cells were counted 1.0 to 1.5 mm to the margin of the optic nerve head. The distribution of EPOR was examined, the primary antibodies of goat anti-EPOR (1:50) and mouse anti-Thy-1 (1:50) were stained overnight. These were then washed thrice with PBS and the secondary antibody of Alexa Fluor 488-labeled donkey anti-goat IgG (1:200) was incubated for 30 minutes. After the washing, Alexa Fluor 594-labeled goat anti-mouse IgG was reacted and washed three times with PBS. DAPI was used to detect nuclei in fluorescence.

### Statistical analysis

All data was expressed as means ± standard deviation. The data was analyzed using t-test and one-way analysis of variance (ANOVAs) using excel or SPSS v12.0 software (IBM, USA). Differences with P-values less than 0.05 were considered statistically significant.

## Results

### EPO has neuroprotective function against NMDA on RGCs after 7 days

#### EPO dose-response by retinal flat mount analysis

The retrograde RGC labeling was performed with FluoroGold (FG) and applied to the surface of the optic nerve. The survival rate of RGC was analyzed using retinal flat mount analysis and individually rescued with the following different dose of EPO (0, 10, 50, or 250 ng) (Table 1). After seven days, NMDA-mediated neurotoxicity was effective in triggering RGC death and the survival rates of total, medium, and small RGCs were reduced to 57.85 ± 5.69% (p < 0.001), 62.72 ± 6.83% (p < 0.001), and 52.05 ± 7.28% (p < 0.001) (n = 22). NMDA-mediated neurotoxicity reduced the survival rate of RGC compared to the control groups (which had a 100% survival rate). Large RGCs had a better survival rate due to a low response rate from NMDA-mediated neurotoxicity. Under 10 ng EPO treatment, the survival rate of RGC after NMDA-mediated neurotoxicity was not significantly improved. Under 50 ng of EPO promoted a significantly higher survival rate of the total, medium, and small RGCs after NMDA-mediated neurotoxicity, which were 84.09 ± 8.04% (p < 0.01), 104.53 ± 9.64% (p < 0.001), and 76.22 ± 12.79% (p < 0.01) (n = 7) of the non-EPO treatment, respectively. When the dosage was titrated to 250 ng EPO, it only produced a slightly higher neuroprotective function for medium RGCs and the survival rate was 82.87 ± 12.46% (p < 0.05) (n = 7) compared to the NMDA-treated 50 ng EPO treatment. However, this was not statistically significant. The large RGCs are less responsive to co-administration of NMDA and EPO. (p > 0.05). Overall, 50 ng EPO had a better neuroprotective function on medium and small RGCs, where there was an overall increase in the survival rate of RCG.

**Table 1.** The survival rates of large, medium, and small RGCs was evaluated for different EPO’s dose by retinal flat mount analysis:The survival rates of RGCs were determined as groups (1) Control, (2) NMDA-mediated (80 nmoles NMDA), and (3-5) EPO-treated groups (80 nmoles NMDA + 10, 50, 250 ng EPO).

|  | Survival rate (%) |  |  |  |
| --- | --- | --- | --- | --- |
|  | Total RGCs | Large RGCs | Medium RGCs | Small RGCs |
| (1) Control (n=8) | 100.00±1.84 | 100.00±11.98 | 100.00±2.21 | 100.00±1.97 |
| (2) 80 nmole NMDA (n=22) | 57.85±5.69*** | 83.17±9.83 | 62.72±6.83*** | 52.05±7.28*** |
| (3) 80 nmole NMDA + 10 ng EPO (n=8) | 52.22±6.18 | 91.83±8.38 | 71.54±12.68 | 39.40±15.75 |
| (4) 80 nmole NMDA + 50 ng EPO (n=8) | 84.09±8.04§ | 92.31±9.38 | 104.53±9.64¶ | 76.22±12.79§ |
| (5) 80 nmole NMDA + 250 ng EPO (n=8) | 68.98±7.62 | 85.58±8.99 | 82.87±12.46# | 61.80±14.73 |

The survival rates of RGCs were normalized with control group. The results were statistically analyzed by the Student's t-test. (\* p < 0.05, \*\* p < 0.01 and \*\*\* p < 0.001 compared with control group;

#### EPO Pre/Co/Post-Treatment by retinal flat mount analysis

The neuroprotective function of EPO against NMDA-mediated neurotoxicity pretreatment, co-treatment, and post-treatment were evaluated (Fig 1). The survival rates of total, large, medium, and small RGCs were evaluated by flat mount analysis in NMDA-mediated neurotoxicity at doses of 80 nmoles. The administration of 50 ng EPO 4 or 8 hours pre- or post-NMDA induction was conducted. The co-administration of NMDA and EPO was conducted as the control group (100% survival rate, normalized).

For the pretreatment, EPO was applied 8 hours before NMDA induction. The survival rate of total, large, medium, and small RGCs were 65.77 ± 11.65%, 84.23 ± 13.85%, 61.37 ± 21.97%, and 63.84 ± 25.16% (n = 6), respectively. When the EPO was pretreated 4 hours before NMDA induction, the survival rates of total, large, medium, and small RGCs were 86.00 ± 8.13%, 108.13 ± 6.54%, 84.90 ± 8.45%, and 81.92 ± 14.58% (n = 8), respectively. The shorter pretreatment time (4 hours) had a slightly higher survival rate than the earlier EPO-pretreated time treatment (8 hours), especially on medium RGCs. However, this difference was not statistically significant.

**Fig 1.**
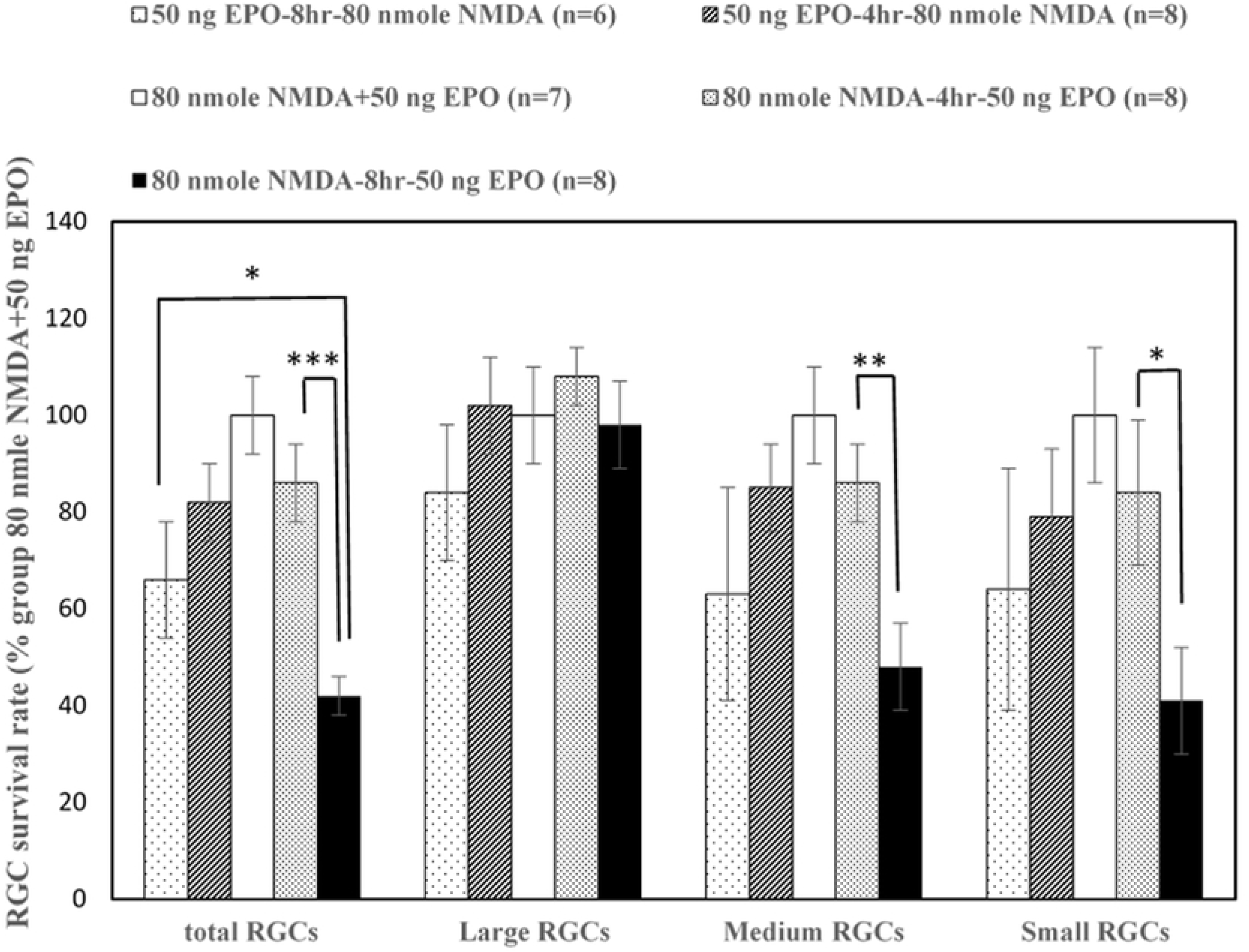
The survival rates of RGCs was determined at different administrating time of EPO by retinal flat mount analysis: The dose of 50 ng EPO was administered pre/co/post injecting intravitreally with 80 nmoles NDMA. The survival rates were normalized with the EPO co-treated groups (80 nmoles NMDA + 50 ng EPO), which is setup for 100% survival rate of RGCs. The survival rates of total, large, medium and small RGCs were determined under different administrating time (pre-treated: 50 ng EPO-4hr-80 nmoles NMDA and 50 ng EPO-8hr-80 nmoles; post-treated: 80 nmoles-4hr-50 ng EPO and 80 nmoles-8hr-50 ng EPO) after 7 days. The co-treated and earlier post-treated groups (0 and 4 hours) had a higher survival rate as compare with the late post-treated group (8 hours) (* p < 0.05; ** p < 0.01; ***p < 0.001 compared with 80 nmoles NMDA-8 hr-50 ng EPO group) (n =6– 8).

For the post-treatment, EPO was applied 4 hours after NMDA induction, the survival rates of total, large, medium, and small RGCs were 82.11 ± 7.85%, 102.34 ± 9.85%, 83.87 ± 9.18, and 77.18 ± 13.62% (n = 8), respectively. When the EPO was applied 8 hours after NMDA induction, the survival rates of total, large, medium, and small RGCs were 41.93 ± 3.84%, 97.25 ± 8.56%, 46.50 ± 9.23%, and 38.69 ± 10.80% (n = 8), respectively. EPO application 4 hours after NMDA induction resulted in a significantly higher survival rate for total (p<0.001), medium (p < 0.01), and small (p < 0.05) RGCs compared to the EPO post-treatment group at 8 hours. In addition, in the same interval of pretreatment and post-treatment, the EPO-pretreated 8 hours NMDA induction had a higher survival rate for total RGCs (p<0.05) compared to the group for EPO post-treated 8 hours after NMDA induction (Fig 1). When the NMDA and EPO were co-administered at the same time, the survival rates of the total, medium, and small RGCs was at the highest compared with the pre- and post-treatment at 4 or 8 hours. The results also illustrated that the large RGCs had much lower response rates pre- and post-treatment, but had more resistance to NMDA toxicity, as indicated by the retinal flat mount analysis.

#### The neuroprotective roles of EPO were evaluated via flow cytometry

We developed a high-content flow cytometer to evaluate the neuroprotective effect of EPO [30]. Seven days after treatment, the survival rates of small and medium RGCs was measured by FG marker in the co-administration of NMDA and EPO (n = 6) [67.43 ± 7.27% (p < 0.001) and 84.4 ± 10% (p < 0.05) respectively], and was significantly higher than the NMDA only group (n = 6) [37.75 ± 6.02% and 60.01 ± 6.01% respectively]. The survival rates of small and medium RGCs treated by combinations of NMDA and EPO as measured by Thy-1 marker (n = 6) [68.14 ± 7.14% (p < 0.001) and 84.4 ± 10.02% (p < 0.05) respectively] was also significantly higher than the NMDA only group (n = 6) [38.60 ± 5.94% and 62.2 ± 6.61% respectively]. On the other hand, the difference in survival rates of large RGCs was not significant between the NMDA-mediated and the NMDA and EPO combination groups. The results demonstrated that EPO has a significant positive effect on the survival rates of small and medium RGCs (Fig 2A). The health status of RGCs can be evaluated using flow cytometry using cell markers, such as FG and Thy-1. For the co-treatment group of 50 ng EPO with 80 nmoles NMDA, FG intensities increased significantly for the large, medium, and small RGCs (n = 6) [87.14 ± 5.71% (p < 0.05), 89.41 ± 2.87% (p < 0.001), and 95.62 ± 2.26% (p < 0.05) respectively] compared to the NMDA-mediated group (n = 6) [68.78 ± 4.47%, 67.07 ± 1.89%, and 88.83 ± 1.79% respectively]. Concurrently, Thy-1 intensities were significantly increased in large, medium, and small RGCs (n = 6) [88.81 ± 6.06% (p < 0.05), 88.93 ± 2.95% (p < 0.001), and 96.02 ± 2.31% (p < 0.05) respectively] compared to the NMDA-mediated group (n = 6) [67.41 ± 4.31%, 66.76 ± 1.76%, and 88.34 ± 1.73% respectively] (Fig 2B). These results demonstrated that EPO may increase the survival rate of small and medium RGCs and overall recover the health of the soma, axon and RGCs.

**Fig 2.**
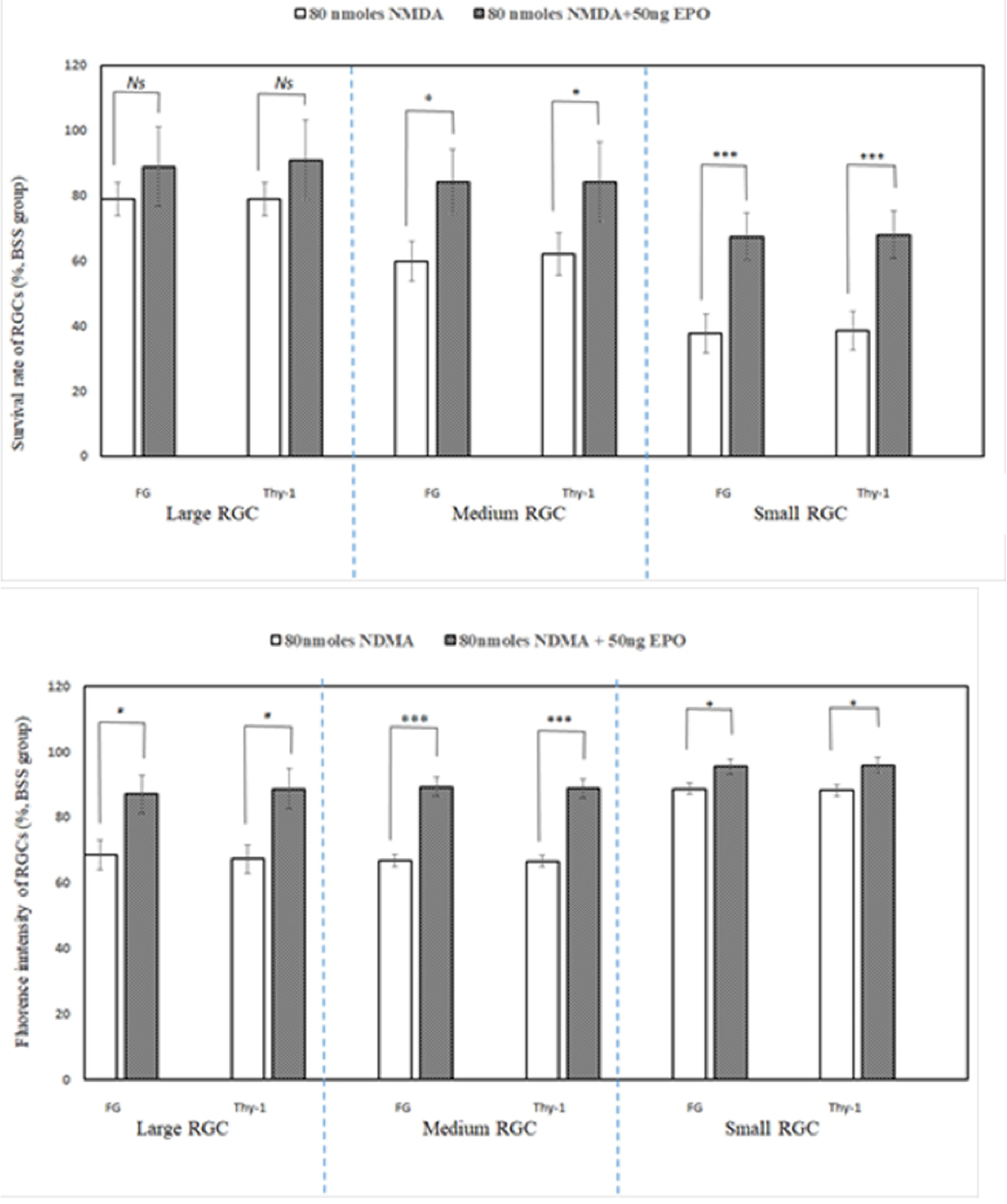
The survival rate and cell health of RGCs were mediated with EPO by flow cytometry (BSS only as 100%): The survival rate of small & medium RGCs were significantly recovered by the treatment of EPO, compared with 80 nmoles NMDA-mediated groups (* p < 0.05; ** p < 0.01; ***p < 0.001) (n = 6) (Upper). EPO-mediated the cell health of all RGCs were evaluated by the intensity of fluorescence (%FG and %Thy-1 intensity) and significantly recovered by the treatment of EPO, compared with 80 nmoles NMDA-mediated groups: (* p < 0.05; ** p < 0.01; ***p < 0.001) (n = 6) (Lower).

### The time course trials of EPO recovered NMDA damage in RGCs

#### EPO rescued the survival rate of RGCs

TUNEL assay is a method used for detecting apoptotic DNA fragmentation. It is widely used to identify and quantify apoptotic cells. In the results of the TUNEL assay in Wistar rats (Fig. 3), the TUNEL-positive cell number in GCL for the BSS-only group (i.e., control treatment) were 15.61 ± 2.71%, 22.29 ± 3.13%, 22.27 ± 2.14%, and 11.03 ± 2.45% at 6, 30, 66, and 90 hours respectively. The TUNEL-positive cell number in GCL were 33.45 ± 4.28% (p < 0.01, n = 4), 42.85 ± 2.07% (p < 0.01, n = 4), 41.62 ± 2.88% (p < 0.01, n = 4), and 37.68 ± 3.39% (p < 0.001, n = 4) respectively, under 80 nmoles NMDA induction compared to the control group. The TUNEL-positive cell number were 30.25 ± 4.05% (p < 0.05), 20.89 ± 2.77% (p < 0.001), and 14.96 ± 3.49% (p < 0.01) at 30, 66, and 90 hours respectively under 50 ng EPO and 80 nmoles NMDA compared to the NMDA induced group. The results demonstrated that EPO rescued the NMDA-mediated neurotoxic cells.

**Fig 3.**
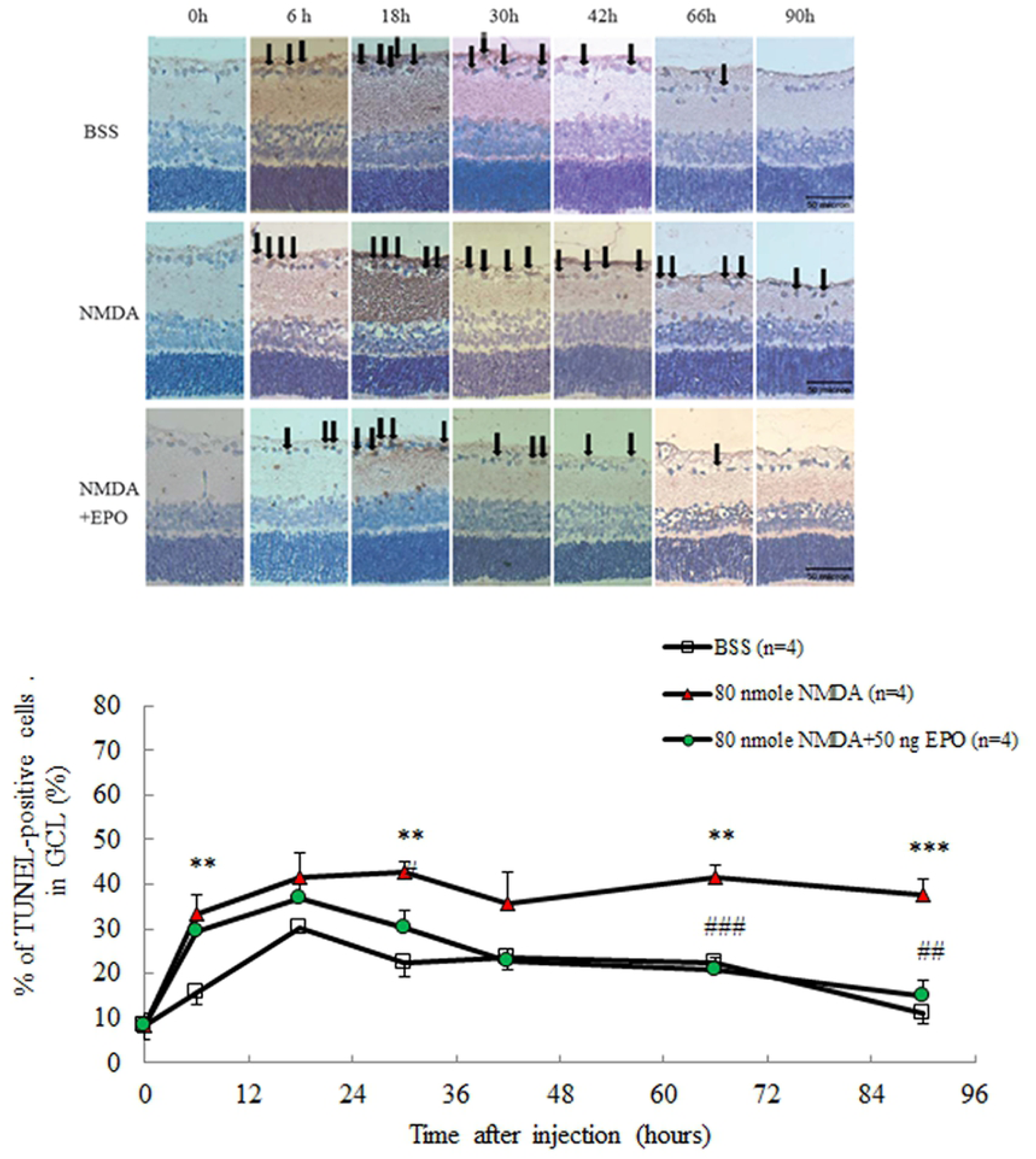
TUNEL analysis were obtained at different analyzing time on Wistar rats: TUNEL analysis at 0, 6, 18, 30, 47, 66, and 90 hrs after receiving the intravitreal injection of (1) 2 μL BSS only, (2) 80 nmoles NMDA, and (3) 80 nmoles NMDA + 50 ng EPO. The positive cell numbers were represented as mean ± SEM (n = 4) (Lower), compared to BSS only (*p < 0.05; **p < 0.01; ***p < 0.001) and compared with NMDA-mediated group (#p < 0.05; ##p < 0.01; ###p < 0.001), Bar = 50 μm.

#### EPO significantly increased the thickness of IPL and the cell number in GCL

The ganglion IPL consists of synaptic connections between the axons of bipolar cells and dendrites of ganglion cells. Recent studies have shown that segmented IPL thickness was associated with glaucomatous damage [12, 42]. In this study, the histological results stained with hematoxylin and eosin (H&E) were examined for the neuroprotective effect of EPO. At the times intervals of 0, 6, 18, 30, 42, 66, and 90 hours, the RGCs counts in the GCL and the ratio of IPL/outer nuclear layer (ONL) were measured (Fig. 4). The results showed that EPO could protect the cells in the GCL and decrease the damage of retinal ganglion dendritic cell and bipolar cell axon induced by NMDA. After 42, 66, and 90 hours after intravitreal injection with 80 nmoles NMDA, the RGCs counts were significantly decreased by NMDA induction compared to the control [85.75 ± 5.89 vs. 61 ± 4.29 (p < 0.01, n = 4), 88.5 ± 5.39 vs. 55.75 ± 6.24 (p < 0.01, n = 4), and 84 ± 6.09 vs. 50 ± 5.1 cells/mm (p < 0.01, n = 4)]. The thickness ratio of IPL/ONL also decreased significantly compared to the control group [0.92 ± 0.07 vs. 0.69 ± 0.06 (p < 0.01, n = 4), 0.91 ± 0.03 vs. 0.58 ± 0.04 (p < 0.001, n = 4), and 0.87 ± 0.07 vs 0.53 ± 0.06 (p < 0.01, n = 4), all respectively]. However, after 42, 66, and 90 hours of intravitreal injection with 80 nmoles NMDA and 50 ng EPO, the RGCs counts were increased significantly to 82 ± 6.98 (p < 0.05, n = 4), 92.5 ± 4.01 (p < 0.001, n = 4), and 82.25 ± 4.25 cells/mm (p < 0.01, n = 4), respectively. The thickness ratio of IPL/ONL also increased significantly to 0.76 ± 0.03 (p < 0.001, n = 4) and 0.73 ± 0.02 (p < 0.05, n = 4) after 66 and 90 hours, respectively.

**Fig 4.**
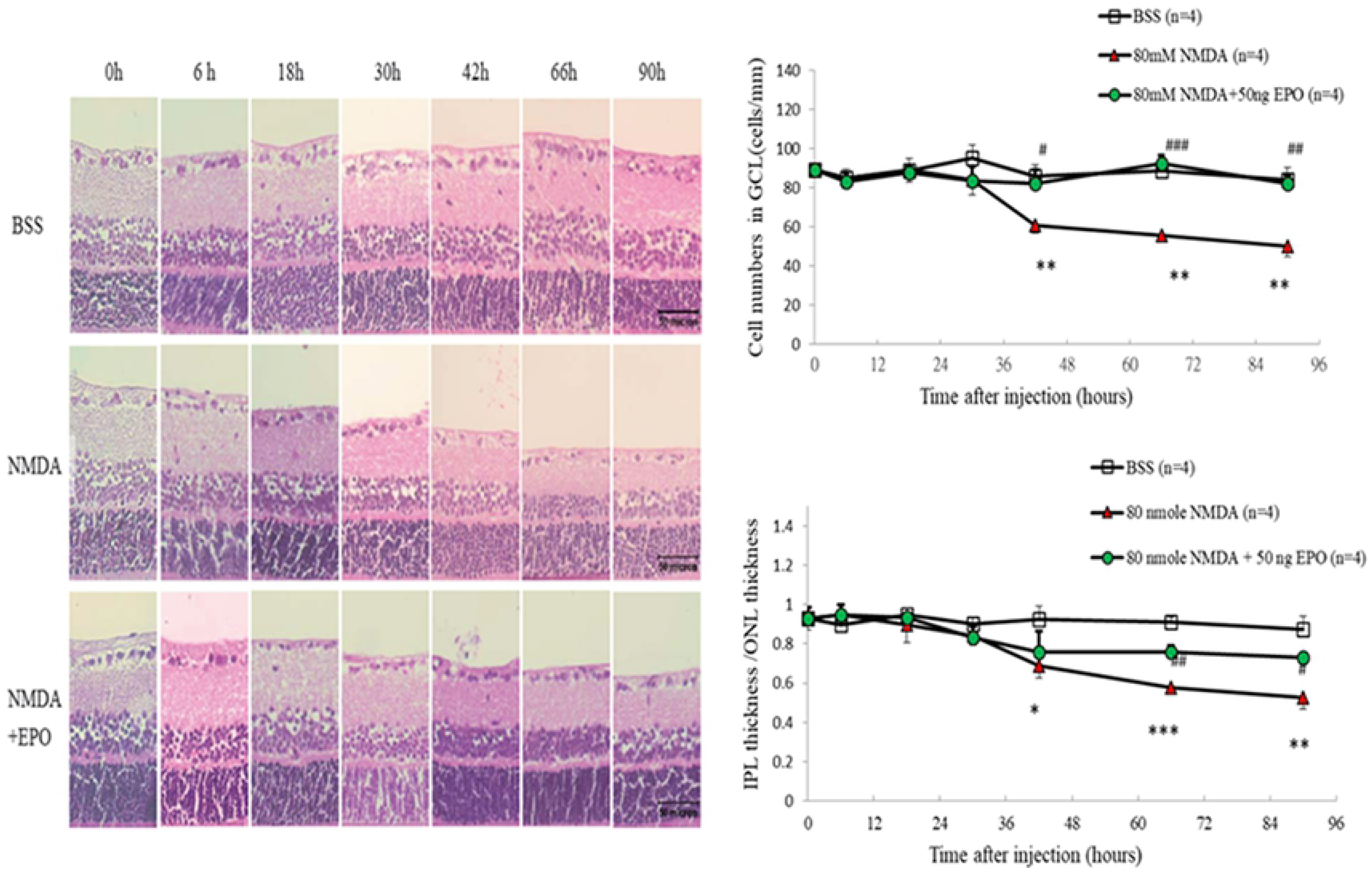
The positive RGC in IPL and the ratio of IPL/ONL were obtained at different analyzing time on Wistar rats: The positive RGC in IPL and the ratio of IPL/ONL of Wistar rats were obtained at 0, 6, 18, 30, 47, 66, and 90 hrs after receiving the intravitreal injection of (1) 2 μL BSS only, (2) 80 nmoles NMDA, and (3) 80 nmoles NMDA + 50 ng EPO; The positive ganglion cell in IPL and the ratio of IPL/ONL were represented as mean ± SEM (n = 4), compared to baseline levels (*p < 0.05; **p < 0.01; ***p < 0.001) and compared with NMDA-mediated group (#p < 0.05; ##p < 0.01; ###p < 0.001), Bar = 50 μm.

#### EPO protected the RGCs by downregulating apoptotic factors

Immunohistochemical staining using an antibody raised against EPOR protein (green color) revealed an intense immunoreactivity in the RGC layer of the retina. Thy-1 (red color) is a surface glycoprotein uniquely expressed on the RGC of retinas. We found an intense immunoreactivity in the RGC layer of Wistar rats. DAPI (blue color) was used for immunohistofluorescence staining. These staining results demonstrated that almost all RGCs were immunoreactive for EPOR (Fig 5).

**Fig 5.**
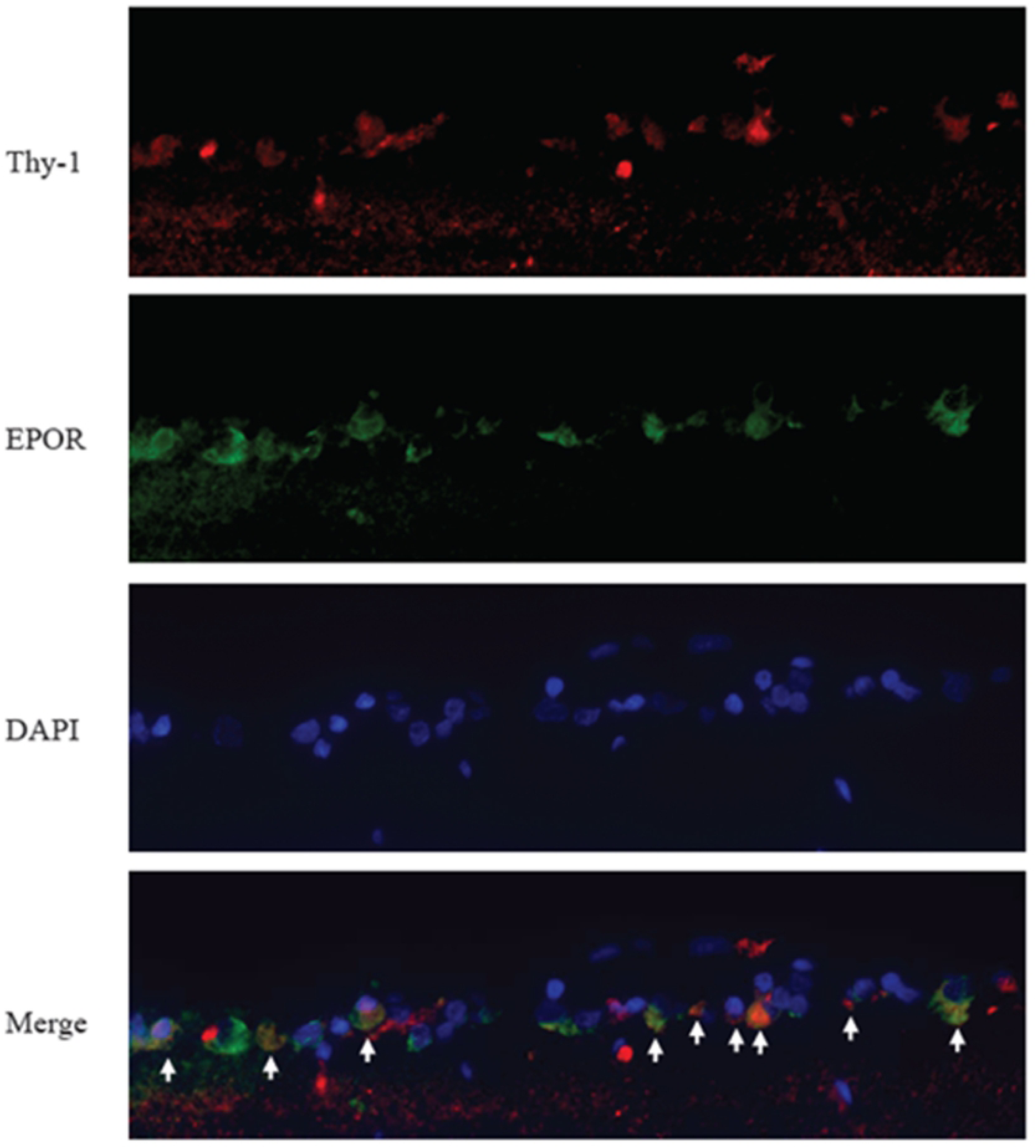
The immunohistofluorescence staining on the ganglion cell layers of Wistar rats : The images showed that Thy-1(red) and EPOR(green) co-expressed with nucleus stained by DAPI (blue).

In order to analyze the signal transduction of RGCs after EPO injection, the apoptotic signals of μ-calpain, caspase 9, and Bax were measured by immunohistofluorescence and immunohistochemistry (S1Fig and Fig 6).

**Fig 6.**
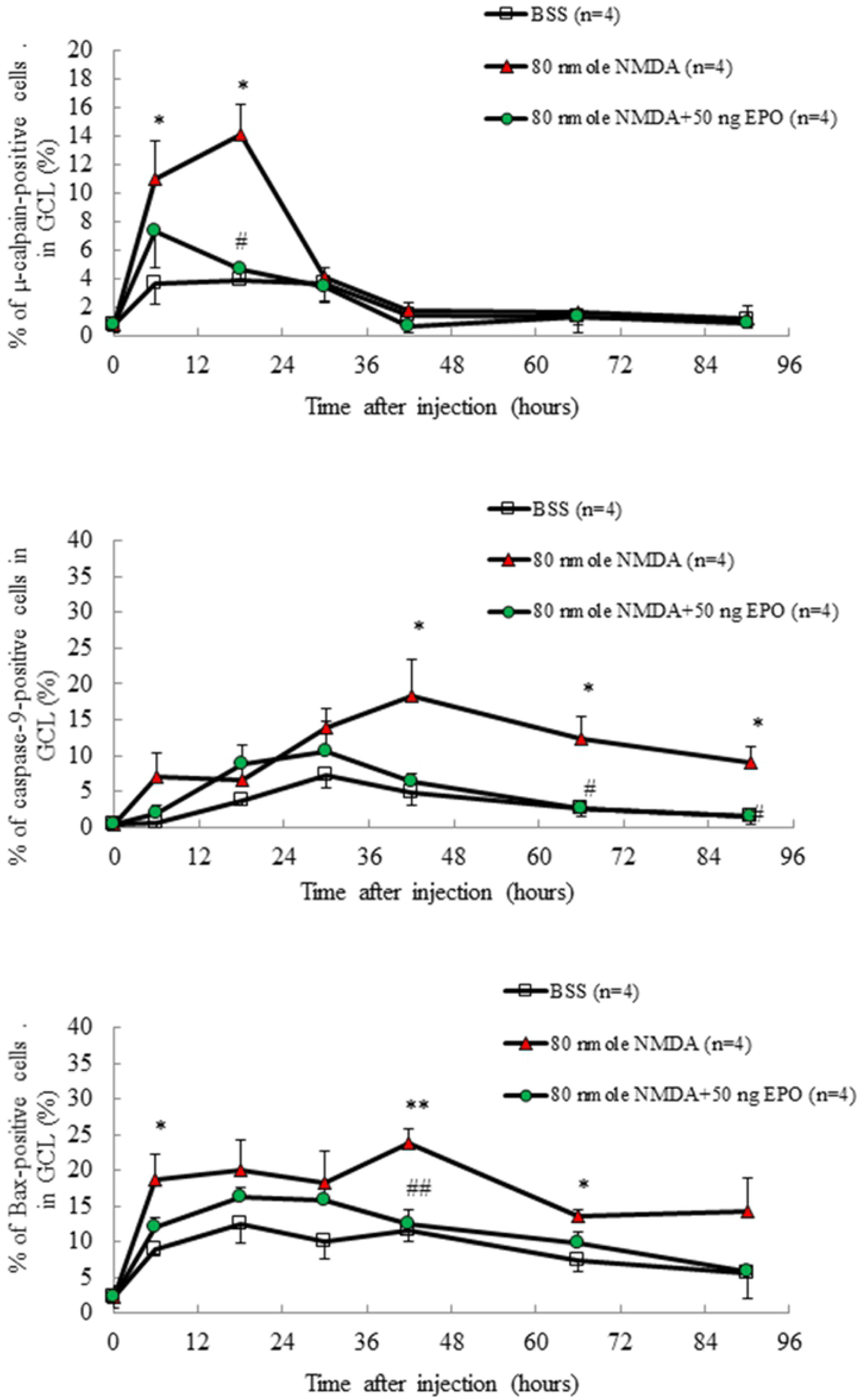
Immunohistochemical staining of µ-caplain-positive, caspase-9-positive, and Bax-positive on ganglion cell layers of Wistar rats: The images were obtained at 0, 6, 18, 30, 47, 66, and 90 hrs after receiving the intravitreal injection of (1) 2 μL BSS only, (2) 80 nmoles NMDA, and (3) 80 nmoles NMDA + 50 ng EPO; the positive cells were represented as mean ± SEM (n = 4), compared with baseline levels (*p < 0.05; **p < 0.01) and compared with NMDA-mediated group (#p < 0.05; ##p < 0.01). (Data derived from S1 Fig.)

For μ-calpain, the percentage of μ-calpain-positive cells in GCL were 3.71 ± 1.52% and 11.01 ± 2.68% (p < 0.05, n = 4) after 6 hours for the control groups (BSS) and the NMDA-mediated groups, respectively. In addition, the percentage of μ-calpain-positive cells in GCL were 3.95 ± 0.15% and 14.10 ± 2.10% (p < 0.05, n = 4) at 18 hours for the control groups (BSS) and the NMDA-mediated groups, respectively. Compared to the NMDA-mediated groups, the percentage of μ-calpain-positive cells in GCL was reduced to 4.64 ± 0.64% (p < 0.05, n = 4) 18 hours after the 80 nmoles NMDA and 50 ng combined EPO treatment.

For caspase 9, the percentage of caspase-positive cells in GCL were 4.90 ± 1.85% and 18.39 ± 4.94% (p < 0.05, n = 4) after 42 hours for the control (BSS) and the NMDA-mediated groups, respectively. The percentage of caspase 9-positive cells in GCL were 2.62 ± 1.22% and 12.23 ± 3.30% (p < 0.05, n = 4) after 66 hours for the control (BSS) and the NMDA-mediated groups, respectively. The percentage of caspase 9-positive cells in GCL were 1.37 ± 0.93% and 8.96 ± 2.29% (p < 0.05, n = 4) after 90 hours for the control and NMDA-mediated groups, respectively. Compared to the NMDA-mediated groups, the percentage of caspase-positive cells in GCL were 2.49 ± 1.02% (p < 0.05) and 1.44 ± 1.11% (p < 0.05) after 66 and 90 hours, respectively, after combining 80 nmoles NMDA and 50 ng EPO treatment.

For Bax, the percentage of Bax-positive cells in GCL were 8.95 ± 0.89% and 18.79 ± 3.39% (p < 0.05, n = 4) after 6 hours for the control (BSS) and NMDA-mediated groups, respectively. The percentage of Bax-positive cells in GCL were 11.58 ± 1.47% and 23.79 ± 2.00% (p < 0.01, n = 4) after 42 hours for the control and NMDA-mediated groups, respectively. The percentage of Bax-positive cells in GCL were 7.38 ± 1.54% and 13.58 ± 0.98% (p < 0.05, n = 4) after 66 hours for the control and NMDA-mediated groups, respectively. Compared to the NMDA-mediated groups, the percentage of Bax-positive cells in GCL was 12.46 ± 2.00% (p <0 .01) 42 hours after combining 80 nmoles NMDA and 50 ng EPO treatment. The present study revealed that erythropoietin (EPO) protects RGCs against NMDA-mediated apoptosis in Wistar rat models. It blocked the generation of pro-apoptotic proteins, such as μ-calpain, Bax, and caspase 9.

## Discussion

Our study has shown that NMDA was able to initialize the apoptosis of RGCs. The EPO treatment can then reverse the NMDA-mediated neurotoxicity on the survival rate of small and medium RGCs by retinal flat mount analysis in an experimental model *in vivo* (Table 1). The large RGCs were more resistant to NMDA-mediated neurotoxicity. Thus, the EPO treatment (10 ng, 50 ng, or 250 ng) did not significantly alter the survival rate. A difference in response between small and large RGCs was also observed in studies of brain-derived neurotrophic factors [44], and in our previous studies *in vitro* [30]. These provide support for the notion that large RGCs are more resistant to the NMDA-mediated toxicity. We also found that the effect of EPO was dose-dependent. Under low dosage (10 ng), there was no evident protective effect across small, medium, or large RGCs. When the dose of EPO increased to 50 ng, the protective effect was significant for the medium and small RGCs. For the EPO dosage of 250 ng, the protective effect on the medium RGCs remained but was absent for the small RGCs. Under the higher EPO dosage (250 ng), its potential neuroprotective effects and underlying mechanism in small RGCs was unclear. Andrews *et al* found that high doses of EPO can have thrombotic toxicities on rats [45]. Therefore, high EPO dosage may have had thrombotic toxicities, causing ischemia and damage to the small RGCs.

We found that with EPO treatment 4 hours after NMDA induction (post-treatment) had higher survival rates for total, medium, and small RGCs compared to the EPO control treatment 8 hours after NMDA induction (Fig. 1). Hence, earlier EPO post-treatment significantly enhanced survival rates, due to the stability and intervention of EPO. In addition, NMDA requires time to induce neurotoxicity on RGC. After 8 hours, the EPO pretreatment group had a higher survival rate for total RGCs than the post-treated group for the post-treatment. This could be due to irreversible, long-lasting NMDA-mediated neurotoxicity on these RGCs after 8 hours, where EPO did not produce a significant effect. Furthermore, NMDA-mediated a significantly higher apoptotic signal (µ-calpain and Bax) than BSS after 6 hours (Fig. 6) and EPO requires a certain amount of time to activate the signal transduction pathway that reduces the apoptosis of RGCs. Therefore, the neuroprotection effect of EPO is more apparent when it is presented prior to the NMDA treatment. However, when we administered NMDA and EPO simultaneously, the survival rates of total, medium, and small RGCs were higher compared to the pretreatment groups after 4 or 8 hours, suggesting that EPO may degrade after several hours. Therefore, EPO was administered too early before the NMDA-mediated neurotoxicity, the concentration of EPO would decline and limiting its neuroprotective effect.

The flow cytometry showed that under NMDA-mediated neurotoxicity, EPO increased the survival rate of small and medium RGCs but not in large RGCs. It also recovered the health of the soma and axon in large, medium, and small RGCs. This result was consistent with our previous retinal flat mount analysis (Fig. 2). This suggests that large RGCs were more resistant to NMDA toxicity. In other words, glaucoma unequally causes more damage to smaller RGCs compared to larger RGCs. This result was also consistent with our previous study using retinal cell culture method *in vitro* [30]. The responses between small and large RGCs were inconsistent across different studies. Mey *et al* showed that cutting down the optic nerve in rats causes the apoptosis of the small RGCs, which was consistent with our study [46]. However, Quigley *et al* showed that chronic glaucoma will selectively damage the large RGCs in monkeys and humans [47–49]. The inconsistency in the results may be attributable to the different experimental conditions and design, such as different animal species and different types of damage to RGCs.

The results of the retinal cross-section analysis by TUNEL assay showed that EPO rescued the NMDA-mediated damaged cells 30 to 90 hours after NMDA injection (Fig. 3). Furthermore, through H&E staining we found that EPO could protect the cells in GCLs and decrease the damage to retinal ganglion dendrites and bipolar cell axon 66 to 90 hours after NMDA toxicity was initiated. These findings were consistent with the flow cytometry and retinal flat mount results, which further confirmed that EPO could not only protect the ganglion cell body, but also protect the ganglion cell dendrites and bipolar cell axon.

The immunohistofluorescence staining *in vivo* demonstrated that EPOR was expressed intensely in the RGCs of Wistar rats. This proved that EPO could act on RGCs via EPOR (Fig. 4). Therefore, EPO may show its neuroprotection effect through binding with EPOR on the ganglion cells, further the activation of the signal transduction pathway in order to inhibit apoptosis of the ganglion cell.

In order to explore the effects of the upregulation of EPO on neuroprotection signal transduction in rat model *in vivo*, the RGCs apoptosis associated factor such as µ-calpain, caspase 9, and Bax were investigated. The percentage of µ-calpain-positive cells were increased significantly in the GCL 6 to 18 hours after NMDA injection, whereas the percentage of Bax-positive cells were significantly increased in the GCL after 6 hours and 42–66 hours after NMDA injection (S1Fig and Fig 6). The percentage of caspase-positive cells were significantly increased in the GCL 42 to 90 hours after NMDA injection. This suggest that during the signal transduction of NMDA-mediated neurotoxicity, µ-calpain was activated first, followed by Bax, and finally caspase 9. The percentage of Bax-positive cells also increased significantly in the GCL 6 hours after NMDA injection. Yet, when the results after 6, 42, and 66 hours were compared, the percentage of Bax-positive cells was the highest after 42 hours, where Bax was activated after µ-calpain. This signal transduction cascade of NMDA-mediated apoptosis was consistent with previous studies [30]. Hartwick *et al* have shown that the activation of the NMDA receptors will cause the death of RGCs, increase the flow of Ca^2+^, and activates µ-calpain in after ∼12 minutes [50]. In glaucoma patients, Wax *et al* found that caspase 8 was activated in the first stage (beginning stage) and caspase 9 was activated in the second stage (decisional stage) during apoptosis of RGCs [51]. EPO rescues the apoptotic pathway to attenuate NMDA-mediated excitotoxic retinal damage. The significant increase of µ-calpain after 6 to 18 hours post-NMDA treatment (Fig. 6) and the poor performance of post-treatment on RGC survival (80 nmoles NMDA, 50 ng EPO) imply that µ-calpain was crucial in determining the survival rate of RGC with EPO treatment. The post-treatment did not produce an effect due to the delayed onset time and the continue rise of µ-calpain. EPO should be administered before the significant rise in µ-calpain, especially for post-treatment application (e.g., 80 nmoles NMDA50 ng EPO after 4 hours) for better outcomes. Schuettauf *et al* found that intravitreal injection of NMDA will cause cell apoptosis through extrinsic and intrinsic pathways [52]. On the extrinsic pathway, procaspase 8 will be activated to caspase 8, which will subsequently activate caspase 3. On the intrinsic pathway, the Bax will migrate to the mitochondria, where it promotes the mitochondria to release cytochrome C. The cytochrome C will activate caspase 9 which will then activate caspase 3. Caspase 3 will further activate caspase-activated DNase, causing damage to the DNA. Our study confirmed that EPO presents the neuroprotection effect by downregulating the activity of µ-calpain, Bax, and caspase 9, which results in the reduction of apoptosis of the ganglion cell through binding EPOR on the ganglion cells. According to our study and previous studies, we postulated the possible pathway of neuroprotection of EPO in the apoptotic pathway (Fig. 7).

**Fig 7.**
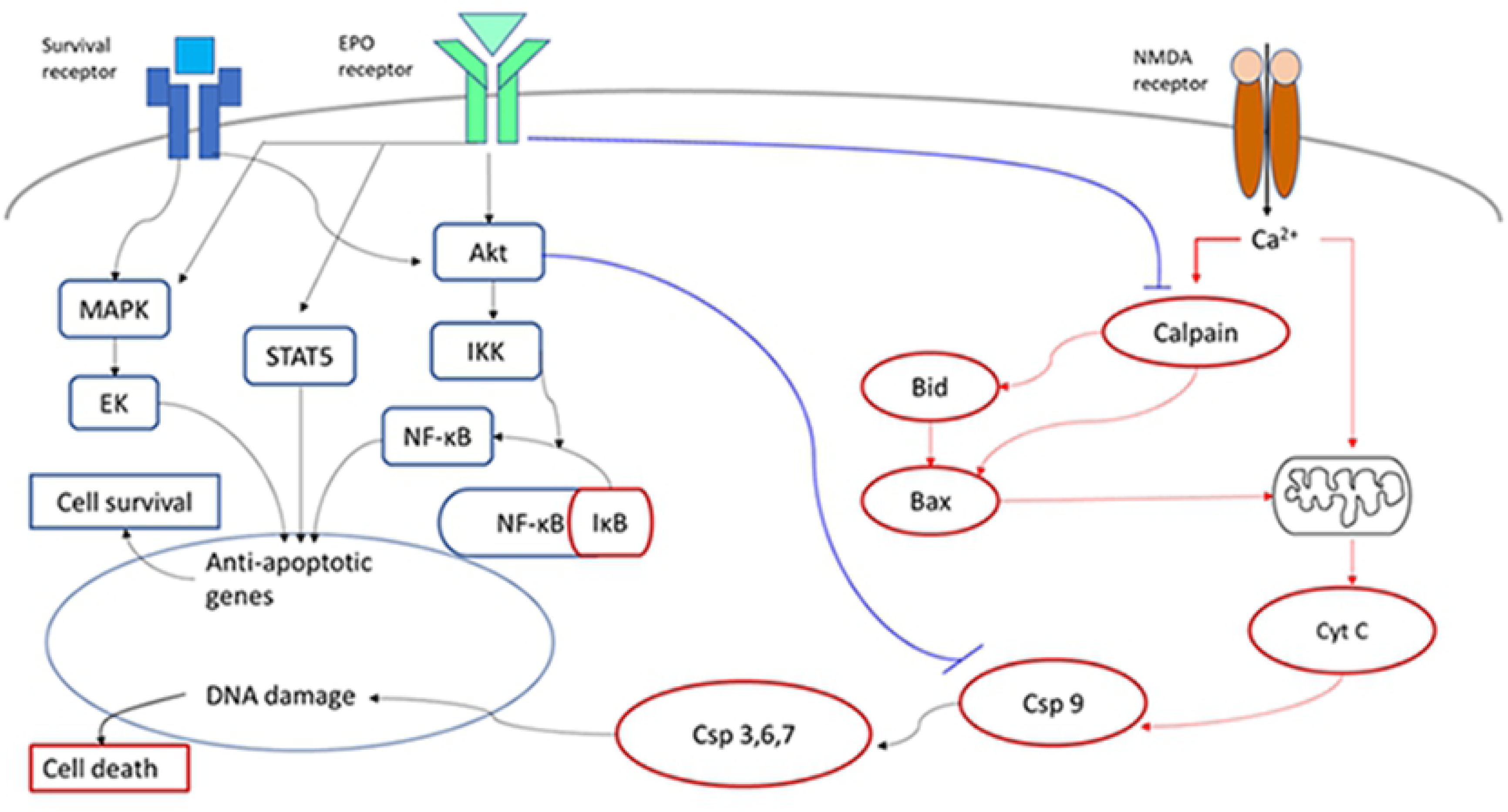
The possible signal transduction pathways: activated by NMDA-mediated neurotoxicity (red line) and downregulated by EPO-mediated neuroprotective (blue line) in the apoptotic pathway.

Our immunohistochemical results at different time intervals revealed that NMDA requires time to induce neurotoxicity on RGC, and that EPO requires time to intervene the apoptosis cascade. Thus, it is imperative to urgently find an efficient delivery method for EPO to recover the apoptosis. Nevertheless, EPO can modulate programed cell death and may be a good potential therapeutic candidate for neuroprotection. This can potentially be used to treat glaucoma.

## Conclusions

Our results demonstrated that NMDA induces neurotoxicity in the majority of small and medium RGCs, and EPO treatment can protect RGC bodies, ganglion cell dendrites, and bipolar cell axon from NMDA-mediated toxicity. In addition, EPO expressed a dose-dependent effect where increasing the EPO dose significantly increases the protective effect in medium and small RGCs. Furthermore, EPO requires a certain amount of time to activate the signal transduction pathway to reduce the apoptosis of the ganglion cell. Thus, the EPO expresses better neuroprotection effect if it is administered earlier once the NMDA toxicity begins, and even before the toxicity is induced. Our findings suggest that there are EPORs on the RGCs. Hence, the EPO may show its neuroprotection effect through binding EPOR on the ganglion cells, which activates the signal transduction pathway to inhibit apoptosis. Furthermore, we observed that in this signal transduction pathway µ-calpain is activated first, followed by Bax, and then caspase 9. We also confirmed that EPO produces the neuroprotection effect by downregulating the activity of µ-calpain, Bax, and caspase 9, and facilitating the reduction of apoptosis in the ganglion cells of Wistar rats. Future studies will focus on the neuroprotective effects of EPO on human ganglion cells to explore its potential application for glaucoma patients.

## Funding

This work was supported by grants from Tri-Service General Hospital (TSGH-C108-120) and Ministry National Defense-Medical Affairs Bureau (MAB-109-032; MAB-109-086).

## Acknowledgments

The authors acknowledge the technical services provided by Instrument Center of National Defense Medical Center.

## Conflicts of Interest

The authors declare no conflict of interest.

## Supporting information

**S1_Fig Immunohistochemical staining of µ-caplain-positive, caspase-9-positive, and Bax-positive on retinal ganglion cell layers of Wistar rats:** Immunohistochemical staining on GCLs of Wistar rats were obtained at 0, 6, 18, 30, 47, 66, and 90 hrs after receiving the intravitreal injection of (1) 2 μL BSS only, (2) 80 nmoles NMDA, and (3) 80 nmoles NMDA + 50 ng EPO; the positive cells were represented as arrows. (a) µ-caplain-positive, (b) caspase-9-positive and (c) Bax-positive cells.

## References

1. Quigley HA, Broman AT. The number of people with glaucoma worldwide in 2010 and 2020. Br J Ophthalmol. 2006; 90(3): 262–7.

2. Bourne RR, Taylor HR, Flaxman SR, Keeffe J, Leasher J, Naidoo K, Pesudovs K, White RA, Wong TY, Resnikoff S, Jonas JB. Number of People Blind or Visually Impaired by Glaucoma Worldwide and in World Regions 1990 - 2010: A Meta-Analysis. PLoS One. 2016; 11(10): e0162229.

3. Yamazaki M, Omodaka K, Takahashi H, Nakazawa T. Estimated retinal ganglion cell counts for assessing a wide range of glaucoma stages, from preperimetric to advanced. Clin Exp Ophthalmol. 2017;45(3):310–313.

4. Chidlow G, Wood JP, Casson RJ. Pharmacological neuroprotection for glaucoma. Drugs. 2007; 67(5):725–59.

5. Fuchs C, Forster V, Balse E, Sahel JA, Picaud S, Tessier LH. Retinal-cell-conditioned medium prevents TNF-alpha-induced apoptosis of purified ganglion cells. Invest Ophthalmol Vis Sci. 2005; 46(8): 2983–91.

6. McKinnon SJ, Schlamp CL, Nickells RW. Mouse models of retinal ganglion cell death and glaucoma. Exp Eye Res. 2009;88(4): 816–24.

7. Quigley HA, McKinnon SJ, Zack DJ, Pease ME, Kerrigan-Baumrind LA, Kerrigan DF, Mitchell RS. Retrograde axonal transport of BDNF in retinal ganglion cells is blocked by acute IOP elevation in rats. Invest Ophthalmol Vis Sci. 2000; 41(11): 3460–6.

8. Pease ME, McKinnon SJ, Quigley HA, Kerrigan-Baumrind LA, Zack DJ. Obstructed axonal transport of BDNF and its receptor TrkB in experimental glaucoma. Invest Ophthalmol Vis Sci. 2000; 41(3): 764–74.

9. Nakazawa T, Nakazawa C, Matsubara A, Noda K, Hisatomi T, She H, Michaud N, Hafezi-Moghadam A, Miller JW, Benowitz LI. Tumor necrosis factor-alpha mediates oligodendrocyte death and delayed retinal ganglion cell loss in a mouse model of glaucoma. J Neurosci. 2006; 26(49): 12633–41.

10. Zhou X, Li F, Kong L, Tomita H, Li C, Cao W. Involvement of inflammation, degradation, and apoptosis in a mouse model of glaucoma. J Biol Chem. 2005; 280(35): 31240–8.

11. Casson RJ. Possible role of excitotoxicity in the pathogenesis of glaucoma. Clin Exp Ophthalmol. 2006; 34(1): 54–63.

12. Zivkovic M, Dayanir V, Zlatanovic M, Zlatanovic G, Jaksic V, Jovanovic P, Radenkovic M, Djordjevic-Jocic J, Stankovic-Babic G, Jovanovic S. Ganglion Cell-Inner Plexiform Layer Thickness in Different Glaucoma Stages Measured by Optical Coherence Tomography. Ophthalmic Res. 2018;59(3):148–154.

13. Newhouse I, Lucas B. The toxic effect of sodium L-glutamate on the inner layers of the retina. AMA Arch Ophthalmol. 1957; 58(2): 193–201.

14. Choi DW. Glutamate neurotoxicity and diseases of the nervous system. Neuron. 1988; 1(8): 623–34.

15. Zhou X, Hollern D, Liao J, Andrechek E, Wang H. NMDA receptor-mediated excitotoxicity depends on the coactivation of synaptic and extrasynaptic receptors. Cell Death Dis. 2013; e560.

16. Siliprandi R, Canella R, Carmignoto G, Schiavo N, Zanellato A, Zanoni R, Vantini G. N-methyl-D-aspartate-induced neurotoxicity in the adult rat retina. Vis Neurosci. 1992; 8(6): 567–73.

17. Dreyer EB, Zurakowski D, Schumer RA, Podos SM, Lipton SA. Elevated glutamate levels in the vitreous body of humans and monkeys with glaucoma. Arch Ophthalmol. 1996; 114(3): 299–305.

18. Hirooka K, Miyamoto O, Jinming P, Du Y, Itano T, Baba T, Tokuda M, Shiraga F. Neuroprotective effects of D-allose against retinal ischemia-reperfusion injury. Invest Ophthalmol Vis Sci. 2006; 47(4): 1653–7.

19. Nivison-Smith L, Acosta ML, Misra S, O’Brien BJ, Kalloniatis M. Vinpocetine regulates cation channel permeability of inner retinal neurons in the ischaemic retina. Neurochem Int. 2014; 66: 1–14.

20. Yoles E, Schwartz M. Elevation of intraocular glutamate levels in rats with partial lesion of the optic nerve. Arch Ophthalmol. 1998; 116(7): 906–10.

21. Marti HH, Wenger RH, Rivas LA, Straumann U, Digicaylioglu M, Henn V, Yonekawa Y, Bauer C, Gassmann M. Erythropoietin gene expression in human, monkey and murine brain. Eur J Neurosci. 1996; 8(4): 666–76.

22. Juul SE, Anderson DK, Li Y, Christensen RD. Erythropoietin and erythropoietin receptor in the developing human central nervous system. Pediatr Res. 1998; 43(1): 40–9.

23. Böcker-Meffert S, Rosenstiel P, Röhl C, Warneke N, Held-Feindt J, Sievers J, Lucius R. Erythropoietin and VEGF promote neural outgrowth from retinal explants in postnatal rats. Invest Ophthalmol Vis Sci. 2002; 43(6): 2021–6.

24. Koury MJ, Bondurant MC. Erythropoietin retards DNA breakdown and prevents programmed death in erythroid progenitor cells. Science. 1990; 248(4953): 378–81.

25. Grimm C, Wenzel A, Groszer M, Mayser H, Seeliger M, Samardzija M, Bauer C, Gassmann M, Remé CE. HIF-1-induced erythropoietin in the hypoxic retina protects against light-induced retinal degeneration. Nat Med. 2002; 8(7): 718–24.

26. Junk AK, Mammis A, Savitz SI, Singh M, Roth S, Malhotra S, Rosenbaum PS, Cerami A, Brines M, Rosenbaum DM. Erythropoietin administration protects retinal neurons from acute ischemia-reperfusion injury. Proc Natl Acad Sci U S A. 2002; 99(16): 10659–64.

27. Sakanaka M, Wen TC, Matsuda S, Masuda S, Morishita E, Nagao M, Sasaki R. In vivo evidence that erythropoietin protects neurons from ischemic damage. Proc Natl Acad Sci U S A. 1998; 95(8): 4635–40.

28. Sirén AL, Fratelli M, Brines M, Goemans C, Casagrande S, Lewczuk P, Keenan S, Gleiter C, Pasquali C, Capobianco A, Mennini T, Heumann R, Cerami A, Ehrenreich H, Ghezzi P. Erythropoietin prevents neuronal apoptosis after cerebral ischemia and metabolic stress. Proc Natl Acad Sci U S A. 2001; 98(7): 4044–9.

29. Ehrenreich H, Hasselblatt M, Dembowski C, Cepek L, Lewczuk P, Stiefel M, Rustenbeck HH, Breiter N, Jacob S, Knerlich F, Bohn M, Poser W, Rüther E, Kochen M, Gefeller O, Gleiter C, Wessel TC, De Ryck M, Itri L, Prange H, Cerami A, Brines M, Sirén AL. Erythropoietin therapy for acute stroke is both safe and beneficial. Mol Med. 2002; 8(8): 495–505.

30. Chang ZY, Yeh MK, Chiang CH, Chen YH, Lu DW. Erythropoietin Protects Adult Retinal Ganglion Cells against NMDA-, Trophic Factor Withdrawal-, and TNF-α-Induced Damage. PLoS One. 2013; 8(1): e55291.

31. Ihle JN, Quelle FW, Miura O. Signal transduction through the receptor for erythropoietin. Semin Immunol. 1993; 5: 375–389.

32. Chong ZZ, Kang JQ, Maiese K. Apaf-1, Bcl-xL, cytochrome c, and caspase-9 form the critical elements for cerebral vascular protection by erythropoietin. J Cereb Blood Flow Metab. 2003; 23(3): 320–30.

33. Uddin S, Kottegoda S, Stigger D, Platanias LC, Wickrema A. Activation of the Akt/FKHRL1 pathway mediates the antiapoptotic effects of erythropoietin in primary human erythroid progenitors. Biochem Biophys Res Commun. 2000; 275(1): 16–9.

34. Silva M, Grillot D, Benito A, Richard C, Nuñez G, Fernández-Luna JL. Erythropoietin can promote erythroid progenitor survival by repressing apoptosis through Bcl-XL and Bcl-2. Blood. 1996; 88(5): 1576–82.

35. Digicaylioglu M, Lipton SA. Erythropoietin-mediated neuroprotection involves cross-talk between Jak2 and NF-kappaB signalling cascades. Nature. 2001; 412(6847): 641–7.

36. Celik M, Gökmen N, Erbayraktar S, Akhisaroglu M, Konakc S, Ulukus C, Genc S, Genc K, Sagiroglu E, Cerami A, Brines M. Erythropoietin prevents motor neuron apoptosis and neurologic disability in experimental spinal cord ischemic injury. Proc Natl Acad Sci U S A. 2002; 99(4): 2258–63.

37. Broker LE, Kruyt FA, Giaccone G. Cell death independent of caspases: a review. Clin Cancer Res. 2005;11(9):3155–3162.

38. McKinnon SJ, Lehman DM, Kerrigan-Baumrind LA, Merges CA, Pease ME, Kerrigan DF, Ransom NL, Tahzib NG, Reitsamer HA, Levkovitch-Verbin H, Quigley HA, Zack DJ. Caspase activation and amyloid precursor protein cleavage in rat ocular hypertension. Invest Ophthalmol Vis Sci. 2002; 43(4): 1077–87.

39. Tan Y, Dourdin N, Wu C, De Veyra T, Elce JS, Greer PA. Ubiquitous calpains promote caspase-12 and JNK activation during endoplasmic reticulum stress-induced apoptosis. J Biol Chem. 2006;281(23):16016–16024.

40. Abri Aghdam K, Soltan Sanjari M, Ghasemi Falavarjani K. Erythropoietin in ophthalmology: A literature review. J Curr Ophthalmol. 2016;28(1):5–11.

41. Kim EK, Park HL, Park CK. Segmented inner plexiform layer thickness as a potential biomarker to evaluate open-angle glaucoma: Dendritic degeneration of retinal ganglion cell. PLoS One. 2017; 12(8):e0182404.

42. Hartwick AT. Beyond intraocular pressure: neuroprotective strategies for future glaucoma therapy. Optom Vis Sci. 2001 Feb;78(2):85–94.

43. Chiu K, Lau WM, Yeung SC, Chang RC, and So KF. Retrograde labeling of retinal ganglion cells by application of fluoro-gold on the surface of superior colliculus. J Vis Exp. 2008; 16: 819–820.

44. Kashiwagi F, Kashiwagi K, Iizuka Y, Tsukahara S. Effects of brain-derived neurotrophic factor and neurotrophin-4 on isolated cultured retinal ganglion cells: evaluation by flow cytometry. Invest Ophthalmol Vis Sci. 2000;41(8):2373–7.

45. Andrews DA, Boren BM, Turk JR, Boyce RW, He YD, Hamadeh HK, Mytych DT, Barger TE, Salimi-Moosavi H, Sloey B, Elliott S, McElroy P, Sinclair AM, Shimamoto G, Pyrah IT, Lightfoot-Dunn RM. Dose-related differences in the pharmacodynamic and toxicologic response to a novel hyperglycosylated analog of recombinant human erythropoietin in Sprague-Dawley rats with similarly high hematocrit. Toxicol Pathol. 2014;42(3):524–39.

46. Mey J, Thanos S. Intravitreal injections of neurotrophic factors support the survival of axotomized retinal ganglion cells in adult rats in vivo. Brain Res. 1993 Feb 5;602(2):304–17.

47. Quigley HA, Sanchez RM, Dunkelberger GR, L’Hernault NL, Baginski TA. Chronic glaucoma selectively damages large optic nerve fibers. Invest Ophthalmol Vis Sci. 1987 Jun;28(6):913–20.

48. Quigley HA, Dunkelberger GR, Green WR. Chronic human glaucoma causing selectively greater loss of large optic nerve fibers. Ophthalmology. 1988 Mar;95(3):357–63.

49. Glovinsky Y, Quigley HA, Dunkelberger GR. Retinal ganglion cell loss is size dependent in experimental glaucoma. Invest Ophthalmol Vis Sci. 1991 Mar;32(3):484–91.

50. . Hartwick AT1, Hamilton CM, Baldridge WH. Glutamatergic calcium dynamics and deregulation of rat retinal ganglion cells. J Physiol. 2008 Jul 15;586(14):3425–46.

51. Wax MB, Tezel G. Immunoregulation of retinal ganglion cell fate in glaucoma. Exp Eye Res. 2009 Apr;88(4):825–30.

52. Schuettauf F, Stein T, Choragiewicz TJ, Rejdak R, Bolz S, Zurakowski D, Varde MA, Laties AM, Thaler S. Caspase inhibitors protect against NMDA-mediated retinal ganglion cell death. Clin Exp Ophthalmol. 2011 Aug;39(6):545–54

